# Zebra finches produce intralaryngeal laminar flow whistles during panting

**DOI:** 10.1101/2025.02.14.638254

**Authors:** Tommi Anttonen, Hugo Loning, Freja M. Felbo, Jakob Christensen-Dalsgaard, Simon C. Griffith, Marc Naguib, Coen P.H. Elemans

## Abstract

Birds and mammals converged upon the same physical mechanism of vocal fold vibration to produce their broad range of voiced sounds critical to communication^1^. The frequency range of vocal fold vibration is limited per species by biophysical constraints to 3-4 octaves^2^. However, recent work reported vocalizations in zebra finches with apparent fundamental frequencies of 7-11 kHz^3^ that far exceed the range of regular calls and song (0.5-1.5 kHz)^4,5^. These “heat” or “incubation” calls are suggested to have close-range communicative relevance in the global temperature rise context^3,6^, but their acoustics are poorly described and by what biophysical mechanism they are produced remains unknown. We recorded heat calls in adult zebra finches *in vivo* and show they are extremely soft, frequency-modulated calls with source levels of 13.9 ± 3.3 dB SPL at one meter with dominant frequencies of 6.8 ± 0.6 kHz. Through a series of *in vitro* experiments, we establish that these calls are aerodynamic whistles produced inside the avian larynx, not syrinx, during inspiration. Respiratory air flow during whistle production is an order of magnitude higher than song and consistent with thermal panting for evaporative cooling^6,7^. Laryngeal geometry and dimensional flow analysis suggest that these whistles are laminar flow whistles that occur when a flow boundary layer is in a transition phase from laminar to turbulent flows^8,9^. Birds, like some rodents^10-12^, are thus able to produce both voiced sounds and aerodynamical whistles in their vocal tract.

## Results and Discussion

We could readily induce heat calls in adult zebra finches by placing them in a temperature-controlled room at 40°C (See **Methods**). The frequency contours of heat calls were typically chevron-shaped with a dominant frequency component at 6.8 ± 0.6 kHz (N=4 animals) (**Fig. 1A-C**), consistent with earlier qualitative descriptions^3,13^. An integer multiple frequency of the dominant frequency was sometimes present (**Fig. 1AB**). Heat calls were very soft with mean source levels of only 13.9 ± 3.3 dB re. 20 µPa at 1 m (N=4) and often nearly masked by background noise levels (**Fig. 1E**). At 10 cm the received level was 33.9 ± 3.3 dB re. 20 µPa (N=4).

**Figure 1.**
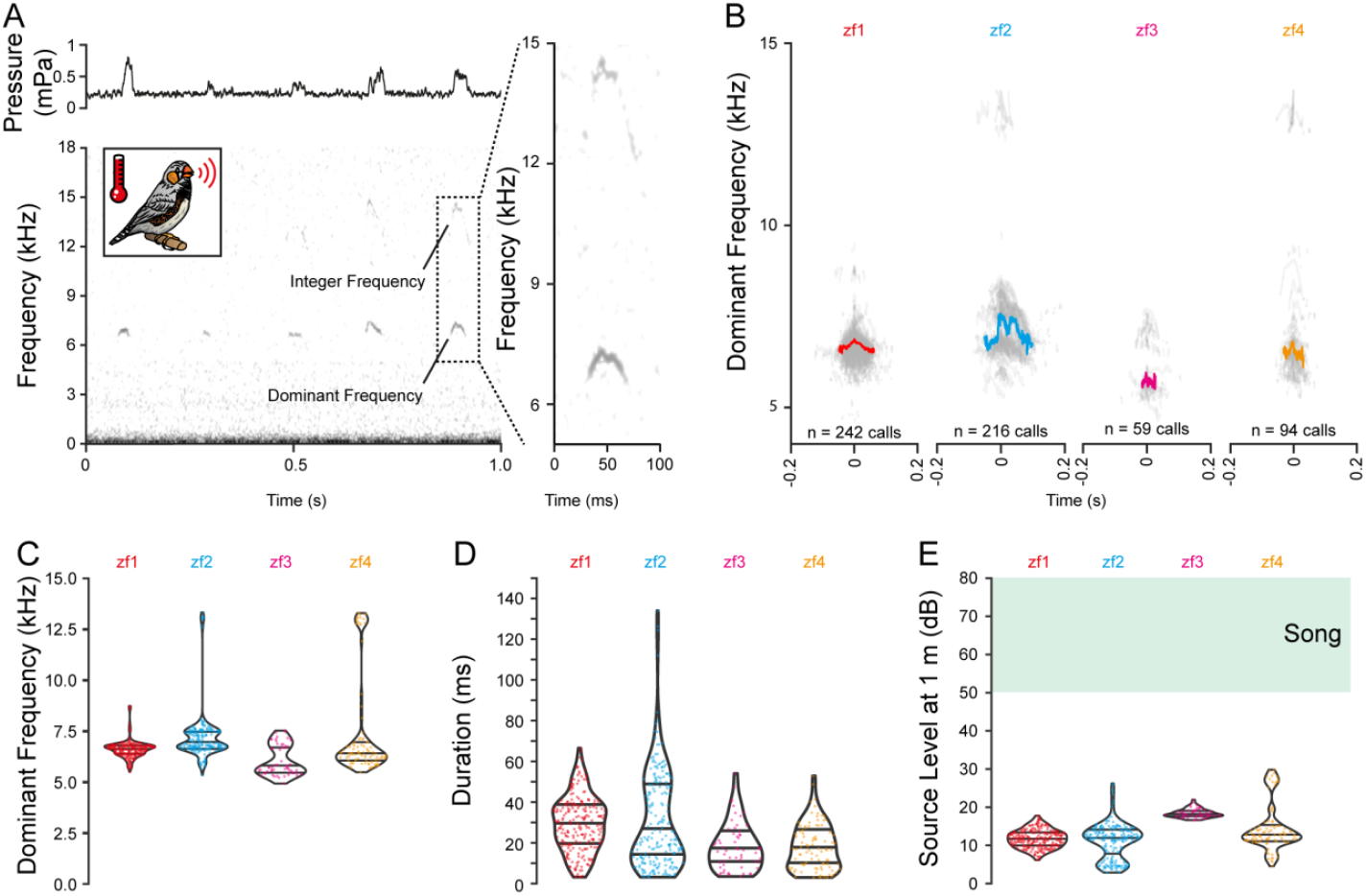
Zebra finch heat calls are extremely soft and high-frequency sounds. (**A**) Representative oscillogram, spectrogram, and a highlighted example (*dashed box*) of heat calls *in vivo*. (**B)** Superimposed frequency traces of all isolated calls (gray) and 90% mean (color) from four zebra finches (*zf1-4*). (**C**) Mean dominant frequency, (**D**) duration and (**E**) source level (mean root-mean-square) of heat calls. Song source level of 50-80 dB is 35 - 50 dB louder^19-22^.

Interestingly, our data strongly suggest that these calls are below the hearing detection threshold of zebra finches. The auditory sensitivity of adult zebra finches at 6-8 kHz ranges from 70 dB SPL (using behavioral test^14^) to 85 dB SPL (using auditory brainstem response tests^15^). Because acoustic pressure does not increase when getting closer to the source in the interference near field, the pressure at 10 cm distance is comparable to sound pressure at the source^16^. Thus, an acoustic pressure of 34 dB SPL at 10 cm is ∼ 35 dB below the behavioral perception limit in adults^14^. In juveniles, hearing sensitivity is strongly age dependent and significantly reduced compared to adults ^15,17^. Embryonic hearing is even less sensitive as hearing gradually develops in altricial birds^15,18^. Thus, embryos, juveniles and adult zebra finches thus cannot detect heat calls as airborne sounds. Our findings therefore raise doubts on the functional significance of zebra finch heat calls in a communicative context^3^.

Previous studies proposed that heat call playbacks at 67 dB SPL at less than 5 cm from egg to speaker stimulate prenatal development in ways that provide an adaptive response to a warming climate^3^. Our findings show that heat call playbacks at 67 dB SPL are at unphysiologically high levels for production of the calls. These earlier studies therefore need to be reproduced at physiologically relevant sound and vibration playback levels and can be best understood by considering the lack of appropriate control groups^13^.

Heat call production seems tightly correlated with respiratory movement of air during breathing, consistent with earlier observations^6,13^, which suggest their origin lies somewhere in the respiratory system between the syrinx and the beak. However, the production of heat calls seems inconsistent with a syringeal sound source for two reasons. First, previous experiments show that vibrating structures in the syrinx produce harmonically rich sounds with a fundamental frequency (*f*o) around 500-1200 Hz^1,4^, which is ∼10 times lower than the lowest frequency component of heat calls (6-8 kHz). Although some males sing “high notes” with a dominant frequency of 4-5 kHz, their rich harmonical content suggest a lower *f*_o_ combined with a 4-5 kHz formant^4^. Second, the chevron-shaped frequency trace (**Fig. 1AB**) coupled to respiratory ventilation strongly suggests that heat call frequency is modulated by air flow as in some rodents ^10,11^. In contrast, frequency-flow coupling is typically weak in flow-induced vocal fold vibration systems such as the syrinx^23^.

To resolve the origin and biophysical mechanism underlying heat call production, we tested whether passive airflow through the isolated upper respiratory tract can induce heat calls *in vitro*. We mimicked expiratory and inspiratory ventilation while measuring air flow and recording produced sounds (**Fig. 2AB**, See Methods). We could successfully induce sounds that closely resemble heat calls with similar frequency of 5.9 ± 2.0 kHz (range: 2.8-8.0 kHz, N=11 animals) and characteristically chevron-shaped frequency modulation (**Fig. 2B**). These sounds occurred consistently during inspiratory flow (i.e. airflow from beak to syrinx) in all tested individuals (N=11 animals), and were only occasionally observed during expiratory flow. Sound frequency was strongly modulated by air volume flow with a slight hysteresis (**Fig. 2C**). These sounds were soft, with source levels of 43.4 ± 7.2 dB re. 20 µPa at 1 m (range: 30.7-53.2 dB, N=11 animals; **Fig. 2D**). The volume flow magnitude during heat calls was 8.9 ± 4.4 ml/s (range: 2.6-27.0 ml/s, N=11 animals; **Fig. 2E**), which is an order of magnitude above quiet respiration (0.8-1.7 ml/s), 3-10 times higher than during singing (1.3-2.0 ml/s), and consistent with flow magnitudes observed only during (thermal) panting (2.4 *-*10 ml/s)^7^. Thus, heat calls are not produced by the syrinx, but somewhere in the upper respiratory tract between the trachea and the beak.

**Figure 2.**
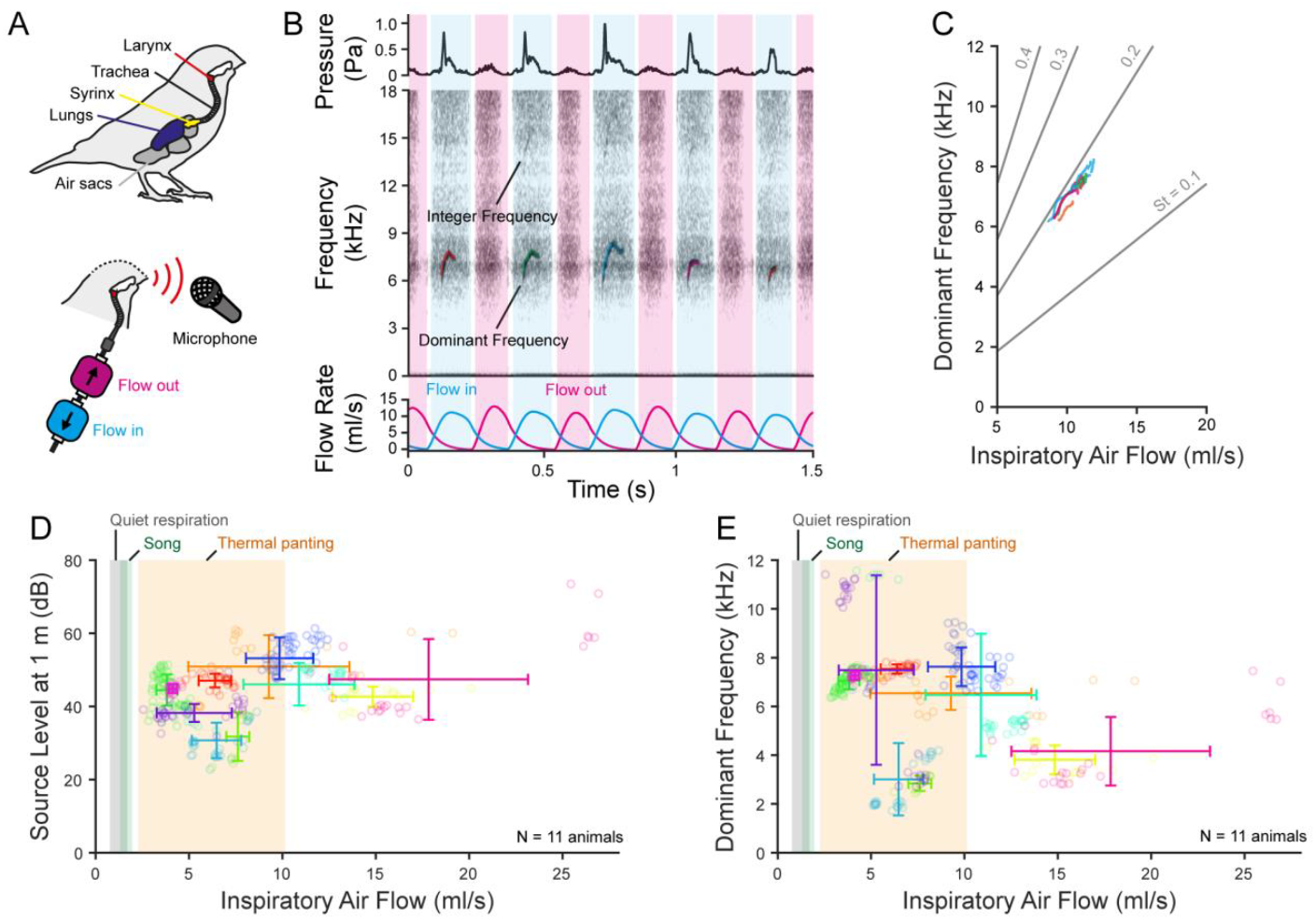
Heat calls are produced in the upper vocal tract during inspiratory flows. (**A**) Schematic of the avian respiratory system with setup to induce heat calls in an isolated larynx *in vitro*. (**B**) Sound amplitude, spectrogram and airflow measurement of a heat call series *in vitro*. Air flow direction is indicated by blue (inspiratory) and purple (expiratory) shaded areas. (**C**) The dominant frequency increases with air flow rate. Strouhal number (St) isolines indicate the relation and range of predicted stable whistles (See **Fig 3F-H**). (**D**) Mean source level and (**E**) mean dominant frequency versus inspiratory flow shows that heat calls are produced during inspiratory flows with magnitudes as seen during thermal panting^7^ or higher.

To more precisely pinpoint the anatomical origin of heat call production, we next systematically explored potential sound-producing structures within the larynx. The strong correlation between flow and sound frequency (**Fig. 2C**) suggests that these calls are aerodynamic whistles, as also found in the rodent larynx^10,11^. Therefore we were particularly alert to flow channel geometry, such as sharp edges and sudden canal narrowing or widening that can cause flow speed increase or flow separation needed for aerodynamic whistles^9^. The avian larynx consists of three cartilages that shape the inner air way geometry; (1) the cricoid is a highly asymmetrical cylinder that dorsally embraces the unpaired (2) procricoid, which supports the bilateral ventrally placed (3) arytenoids (**Fig. 3A-C**)^24,25^. The arytenoids lack vocal folds and are only covered by thin epithelia that outline the glottis, which is open in rest position (**Fig. 3B**). However, we found soft tissues that could exhibit airflow-induced vibration on both the (L1) anterior and (L2) posterior end of the glottis (**Fig. 3D**). On the inside of the flow channel several features could induce aerodynamic whistles. Most notably, (L3) the decrease of tube diameter, (L4) a procricoid ridge protruding into the laryngeal lumen, and (L5) the circular edge of the tube for mounting the excised larynx (**Fig. 3B-D**).

**Figure 3.**
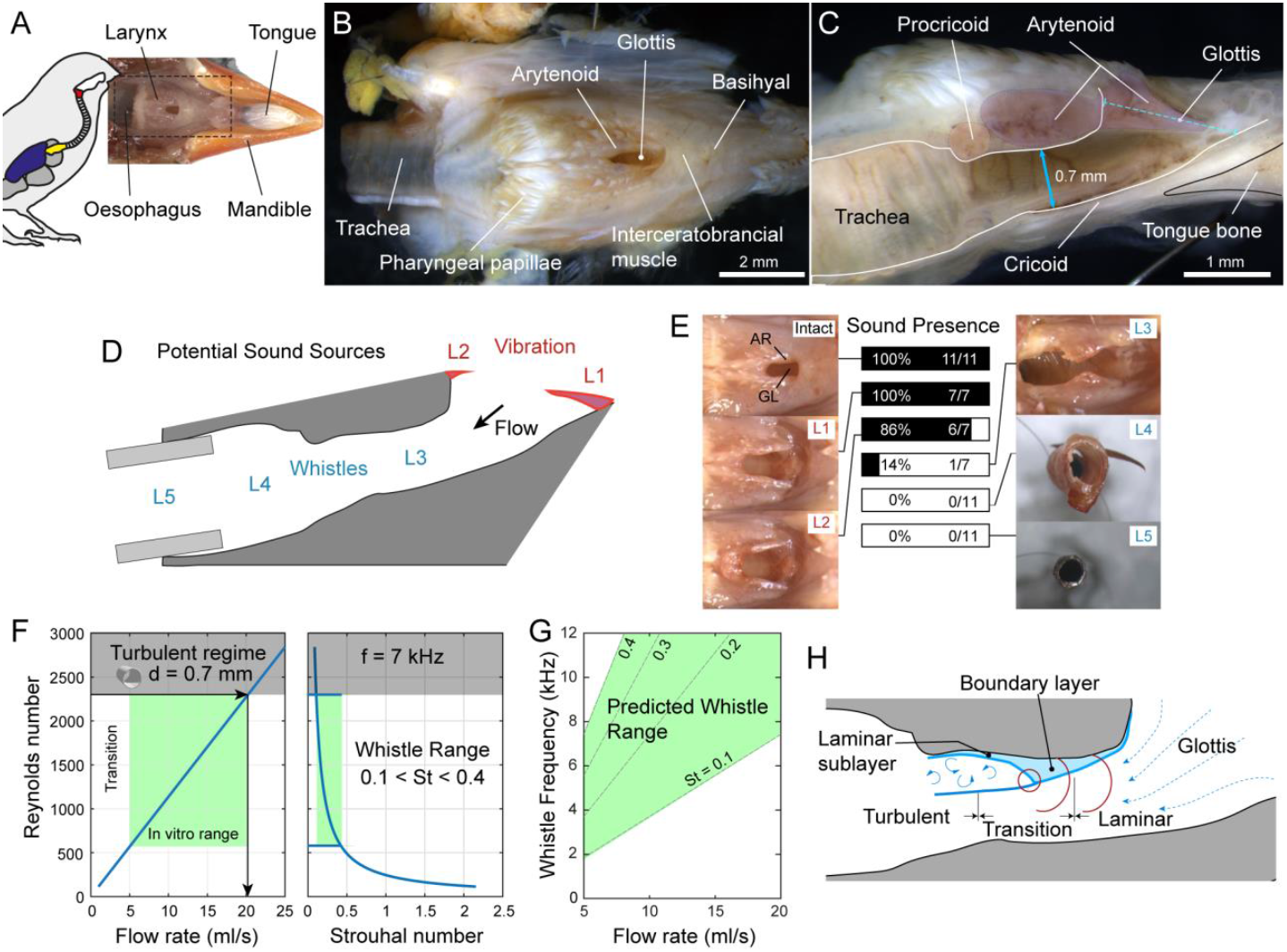
Heat calls are intralaryngeal laminar whistles. (**A**) Overview of larynx position within the pharynx and (**B**) detailed top view and (**C**) medial cross-section of the zebra finch larynx. (**D**) Potential sound sources including vibratory soft tissues on (L1) anterior and (L2) posterior end of the glottis, and whistles where the (L3) flow channel diameter decreases and the procricoid forms a ridge, (L4) trachea and (L5) the tube edge for mounting the excised larynx. (**E**) Systematic surgical removal of potential sounds sources (as in D) shows that removal of the dorsal wall of the flow channel abolishes heat call production. *AR* = Arytenoid; *GL* = Glottis. (**F**) Observed air flow rates (green box) and larynx geometry are consistent with Reynolds and Strouhal numbers for known whistle types. (**G**) Predicted frequencies (*green box*) are consistent with observed data (compare to Fig. 2CE). (**H**) Suggested model for inspiratory whistle production within the avian larynx.

By systematically transecting the identified structures, we found that heat calls are not made by mucosal vibration on either anterior or posterior side of the glottis (**Fig. 3E**). Instead, heat call production ceased in 86% of cases (6 out of 7) when the rigid arytenoid/procricoid was removed (**Fig 3E**). Taken together, our data support the hypothesis that heat calls are aerodynamic whistles that are produced as inspiratory air flow passes through the avian larynx.

Aerodynamic whistles rely on both flow instability and some form of feedback that enhances the instability^9,26-28^. A common excitation mechanism that triggers feedback for most whistles is a flow instability that originates from shear flow, which occurs when the flow speed reaches a certain critical value. If flow speed increases further the flow becomes turbulent which generates white noise (hiss), which is not a whistle. Dimensional analyses are critical to understand favorable flow condition for whistles to occur and particularly relevant are the dimensionless ratio between steady inertial forces to the steady viscous forces (i.e., the Reynolds number *Re*) and the dimensionless ratio between unsteady and steady inertial forces (i.e. the Strouhal number *St*). In a tube, fluid flow transitions from laminar to turbulent at Reynolds numbers around 2300<Re<3000^29^, and thus the flow instability required for whistles predicts that whistles should start at *Re* >2300 (**Fig. 3F**).

Applying dimensional flow analysis to the zebra finch larynx predicts that flow starts transitioning to turbulent at the narrowest point (0.7 mm) in the laryngeal flow channel at 20 ml/s (**Fig. 3C**; See **Methods**). The flow range where we observed whistles in vitro (5-20 ml/s) corresponds to Reynolds numbers of 600<*Re*<2300 and, at a tone of 7 kHz, to Strouhal number of 0.1<*St*<0.4 (**Fig. 3F**). Although Reynolds number approximations are thus below the limit for flow instability in a tube, aerodynamic whistles can occur as low as *Re*>800^28^, which is consistent with our observation. Furthermore, generation of all types of whistles typically occurs within flow conditions with Strouhal number ranging from 0.1 to 0.4^30^, which is also in agreement with our observations. Thus, flow conditions in the larynx support whistle generation and the predicted frequency-flow relationship (**Fig 3G**), is consistent with the observed frequency range (**Fig. 2CE**).

The most common feedback mechanisms in whistle production are fluid or acoustic resonances, which can lead to stable vortex shedding and very loud sounds^28^. Combining the observed air flow rates with larynx geometry, we can exclude the possibility that the mechanism of whistle production is (1) an aeolian tone (because this requires flow over an isolated structure), (2) a pipe tone (because the observed frequencies increase with flow and do not lock), (3) a trailing edge tone (because there is no plate in the flow field) or a (4) resonance-driven wall tone whistle observed in mice and rats^10,11^ (because the flow does not collide with a structure downstream).

Although we cannot exclude the possibility that acoustic resonances play a role in heat whistle production as a feedback mechanism, this seems unlikely. At whistle frequencies of 7 kHz the wavelength is ∼49 mm, which is much larger than any geometric structure in the flow channel. Even a ¼ wavelength resonance would still require a ∼12 mm standing wave. Also, resonance driven whistles are typically rather loud, which is not the case here.

We suggest that avian laryngeal whistles are produced when the boundary layer is in a transition phase from laminar to turbulent flow in the region between arytenoid and procricoid (**Fig 3H**). Boundary layer transitions are very complex and not well understood, but during the transition from laminar to turbulent, pressure fluctuations can cause rapidly changing flow structures that produce tones. To our best knowledge, such so-called laminar flow whistles have been found only to occur along car mirrors^8,9^. In zebra finches laminar flow whistles occur when flow transitions to instable at rates associated only with thermal panting (**Fig. 2E**). The elevated water loss previously attributed to “vocal panting”^6^ can be thus explained by increased respiratory air flow and does not require tissue vibration.

Interestingly, birds were thought to produce whistles until it was established in the last decade that their vocalizations are produced by syringeal vocal fold vibrations^1,31^. Our current findings show that birds, like rodents, can employ both voiced and whistle mechanisms to generate sounds. Particularly, the very high fundamental frequencies produced by e.g. Sharp-beaked ground finches (*Geospiza difficilis*) from the Galapagos islands (*f*_o_ = 10-16 kHz)^32^ or Ecuadorian Hillstar (*Oreotrochilus chimborazo*) hummingbird (f0 = 13.4 kHz)^33^ are interesting candidates. These could be either whistles or vocal fold vibration in higher vocal registers as observed in human classical singers^34^ and toothed whales^35^. Irrespective of what mechanism generates these sounds, the evaluation of their possible communicative value critically requires knowledge and consideration of both sound source level and the hearing capacity of the receiver.

## Funding

This work was supported by Lundbeck Foundation grant to TA and Novo Nordisk Foundation grant NFF20OC0063964 to CPHE.

## Author contributions

TA, FMF and CPHE conceptualized in vitro experiments; HL, SCG, MN conceptualized in vivo experiments, which were conducted by SCG and HL (data for figure 1). TA, JCD and FMF collected data for figures 2 and 3. TA and CPHE analyzed the data. CPHE wrote the draft of the manuscript. All authors contributed to final draft; TA and CPHE acquired funding.

## Declaration of interests

None.

## Methods

### Animals

*In vivo experiments:* Four male zebra finches were used to record heat calls *in vivo*. These males were wild-derived zebra finches housed in outdoor aviaries at Macquarie University. The population was based on wild-type birds that were caught in North-Western New South Wales in 2007 and 2010. The birds were maintained in mixed sex flocks of 30-50 birds.

*In vitro experiments:* Eleven zebra finches (6 males, 5 females) bred at the University of Southern Denmark were used to record heat calls *in vitro*. Because heat call acoustics was not statistically significant difference between males and females^3^, we included both sexes in this study. All *in vitro* experiments were conducted at the University of Southern Denmark.

### *In vivo* recordings of heat calls

A bird cage (17 deep x36 wide x30 high cm) with a single perch was placed in a temperature-controlled room set to 40°C. An acoustic logger (Audiomoth, 48 kHz sampling rate, medium gain) was placed 12 cm above the perch. The acoustic logger was calibrated with a 1 kHz pure tone (generated in Audacity 2.2.2) played from a loudspeaker (Tactix BT081B) placed at the location of the perch and facing the logger. Simultaneously, the sound pressure level (SPL) of the played tone at the location of the logger was measured with a SPL meter (Digitech QM1592, C-weighting, slow response, auto level). Zebra finches were placed individually inside the cage for about one hour and the sounds produced by them were recorded by the acoustic logger.

### *In vitro* recordings of heat calls

Animals were euthanized with an overdose of isoflurane (ScanVet, 1000 mg/g) and weighed. The head was removed at the base of the skull, with 10 mm of the trachea included. The cranium and palate, obscuring the dorsal view of the larynx, were removed at the beak junction. The head was supported by orthodontic tray wax (Kerr, 09246) and the trachea was mounted on a blunted 19G needle and secured with a nylon suture (USP 10-0, ARO surgery). The needle was connected to silicon tubing (Dow Corning, inner diameter 4 mm) which was connected to two serially placed flow sensors (PMF2103V, 0-2 L/min, Posifa, Germany), respectively measuring ingressive and egressive mass flow. A 30 ml plastic syringe was manually moved to allow switching between egressive and ingressive flow conditions. The air temperature was measured continuously at 1.5-2.1 cm from the larynx (National Instruments, NI USB-TC01).

Sound was recorded with a high-sensitivity free-field microphone (G.R.A.S, Type 26HH, 55105) mounted 40-63 mm away from the larynx at a 45-degree angle to avoid air flow interfering with the recording. The microphone was connected to a G.R.A.S Power module (type 12 HF) and the sound was low-pass filtered at 10 kHz (THORLABS, EF120). The microphone sensitivity was calibrated before each experiment (Acoustical Calibrator Type 4231, 94 dB SPL, 1000 Hz). Sound and flow signals were digitized at 60 kHz, 16-bit (National Instruments USB-6259).

To induce heat calls , air was pumped in and out of the intact larynx to mimic the air flow in a panting bird at varying flows until heat calls were recorded. A typical recording was 20 seconds with ∼30 flow reversals. Excessive slime production was observed in freshly euthanized animals. Slime and liquid from the dissection were found in the pharynx and glottis and were removed from the intact larynx before heat calls could be produced. After successfully recording heat calls, the following tissues were cut sequentially with micro-scissors to test what part of the larynx was responsible for heat call production.

Manipulation 1 (L1): Removal of interceratobronchial muscle and the underlying rostral tip of the cricoid

Manipulation 2 (L2): Removal of the soft tissue at the caudal glottis.

Manipulation 3 (L3): Removal of the procricoid and the caudal part of the arytenoids.

Manipulation 4 (L4): A ∼3 mm section of the trachea was left on the mounting tube (control).

Manipulation 5 (L5): Empty mouthing needle (control).

After each manipulation, sound was induced for 10 seconds with ∼30 flow reversals. Additionally, pictures of the larynx were made with a Leica camera (Leica DFC425 C) mounted on top of a Leica MX205A stereomicroscope to record the different manipulations. To avoid the tissue from drying out, drops of PBS (0.01 M) were added regularly and pieces of paper towel soaked in PBS were placed on top of the larynx between recordings. Excess liquid was removed before recordings.

### Acoustic analysis of heat calls

The dominant frequency was extracted using the yin-algorithm by manually optimizing minimal aperiodicity (0.05–0.5) and power (3e^-6^ – 3e^-4^). Source level at 1 m distance of the emitted sound was defined as:

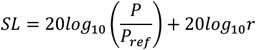

where P is the RMS of sound pressure (Pa), P_ref_ is the reference pressure in air of 20 µPa, and r the distance from the source to the microphone.

### Anatomy

To quantify the physical geometry of the intralaryngeal flow channel, a cross section was performed on bird lbly384. We followed the above protocol and after heat calls had been recorded, we included the control recordings of only trachea and empty needle (see above). Then the entire head was lightly fixed in PFA (4% in PBS, pH 7.4) for 24 hours without any further dissections, so that all of the laryngeal structures remained intact. The larynx was cleaned and cut medially with micro-scissors. Pictures were taken of the larynx submerged in 0.01M PBS with a Leica camera (DFC425 C) mounted on a Leica stereomicroscope (M205A).

### Flow dimensional models

We first analyzed the flow conditions in the larynx based on two dimensionless numbers that provide a first check of fluid flow conditions. The Reynolds number (Re) is the dimensionless ratio between steady inertial forces to the steady viscous forces: 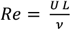, where *U* is flow speed (m/s), *L* a characteristic length (m) and ν the kinematic viscosity of the fluid (m^2^/s). The Strouhal number (*St*) is the dimensionless ratio between unsteady and steady inertial forces: 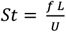, where *f* is frequency (Hz), *U* is flow speed (m/s), and L a characteristic length (m). The smallest tracheal diameter of 0.7 mm was taken as the characteristic length.

## Notes

### Competing Interest Statement

The authors have declared no competing interest.

